# *Klebsiella* Pneumoniae turns more virulent under flow stresses in capillary like microchannels

**DOI:** 10.1101/2023.09.18.558194

**Authors:** Siddhant Jain, Anmol Singh, Nivedita Tiwari, Aparna Naik, Ritika Chatterjee, Dipshikha Chakravortty, Saptarshi Basu

## Abstract

Fluidic habitats are very common to bacterial life, however, very little is known about the effect of the flow stresses on the virulence of the bacteria. In the present work, we conduct microfluidic experiments to understand the consequence of stresses generated by flowing fluid on the bacterial morphology and virulence. We consider *Klebsiella pneumoniae* (*KP*), an ESKAPE pathogen as the model bacteria that are responsible for blood stream infections like bacteremia apart from pneumonia, urinary tract infections and more. We generate four different stress conditions by changing the flow rate and channel geometry subsequently altering the shear rate and stressing time (τ). We observe significant changes in the structural aspects of the stressed bacteria. With an increase in stressing parameters, the viability of the bacterial sample deteriorated. Most importantly, these stressed samples proliferate much more than unstressed samples inside the RAW264.7 murine macrophages. The results shed light on the complex relationship between flow stresses and bacterial virulence. Furthermore, we challenge the bacterial samples with ciprofloxacin to see how they behave under different stress conditions. The present study can be extended to model deadly diseases like bacteremia using organ-on-a-chip technology and help understand bacterial pathogenicity under realistic environments.

**Figure:**
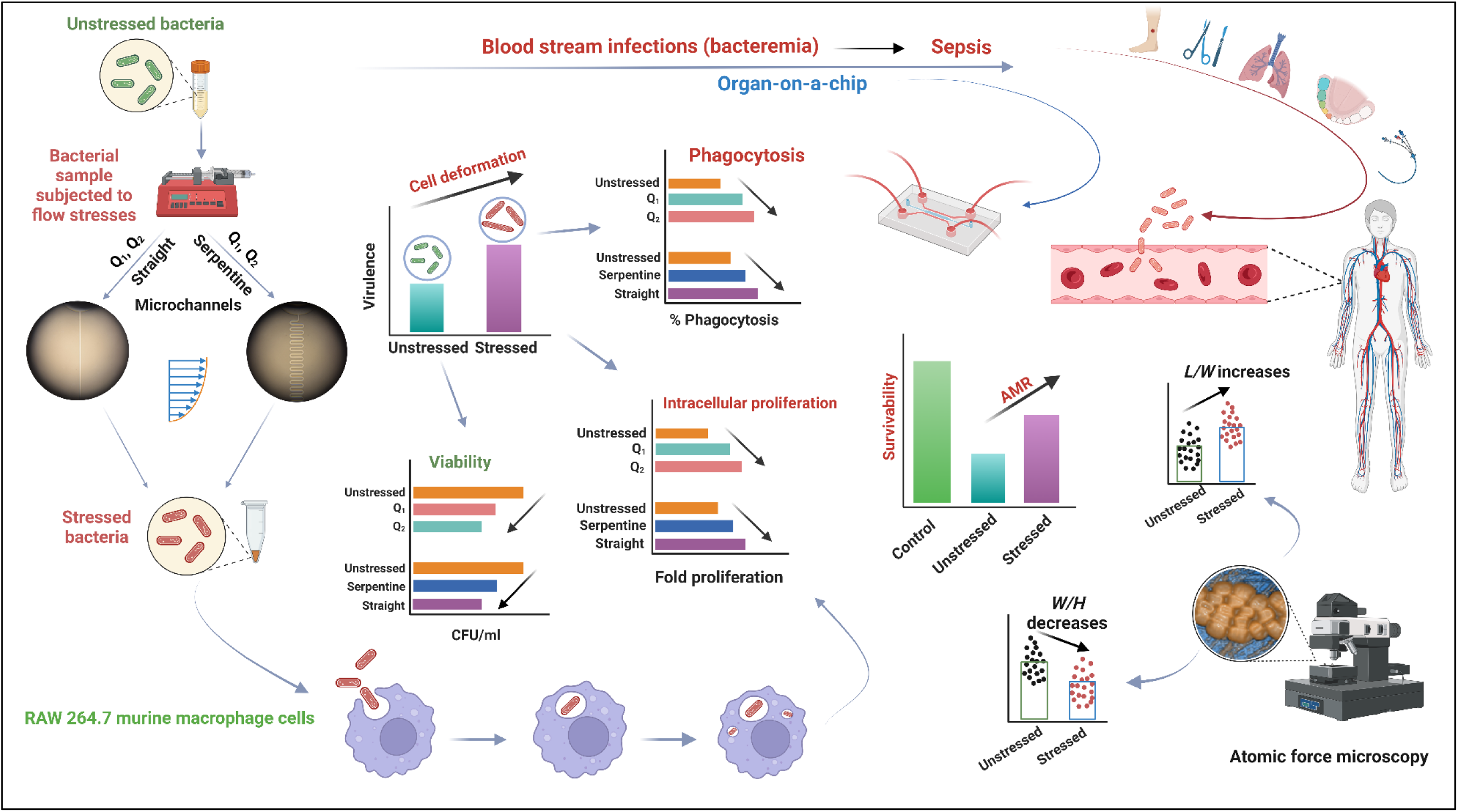
A schematic representation of the present work. Figure created with BioRender (www.biorender.com)

## Introduction

Bacteria constitute 15% of the total global biomass, only behind plants ^1^. They are found in every possible habitat from human body to soils, oceans, snow, etc. Mostly, they flourish under constrained and tortuous environment that plays an influential role on their lifestyle. Being one of the oldest species on Earth, they have evolved significantly and are able to respond to various types of external factors involving electrical, chemical, and mechanical stimuli. Although single-celled, they present a complex case of study involving intricate physio-chemical activities like secretion of extracellular polymeric substances (EPS), constructing protein appendages like pili and fimbriae, formation of biofilms and streamers, quorum sensing, various swimming patterns, taxis and more ^2–7^.

The role of microfluidics has been immense in studying the various aspects of bacterial life. The advancement in the semiconductor industry led to a revolution in the fabrication of small and complex environments for conducting in-vitro studies on these living cells. Few of the important advantages of using microfluidic technologies include easy streamlining of various assays, reduction of reagent volumes (and hence, the cost), better control over biological systems, faster reaction time ^8,9^ etc. The penetration of microfluidics technology in the field of biomedical research has led to various impactful innovations like lab-on-a-chip that have revolutionized point-of-care testing and have solved many laboratory challenges like easy and early detection of diseases^10,11^, modelling of diseases ^12,13^, cell sorting, cell focusing and manipulation ^14–19^, automation and parallelization ^20,21^ to name a few. Using microfluidic techniques Huh et al. ^22^ developed the idea of organ-on-a-chip (OoC) that promises to change the traditional ways of drug development and modelling diseases.

It is well known that when any pathogen infects a host, it is subject to a multitude of adverse conditions. To survive and colonize, the pathogen hunts for energy sources and suitable niches. This process is balanced by the host immune system that continuously checks for such interactions and activities of microorganism. The metabolic activities taking place in our body can serve to change the virulence of the bacteria. For example, changes in temperature, oxygen deprivation, breakdown of short-chain fatty acids and glucose inside our gut act as signals to induce virulence gene expression ^23^. What is less appreciated is the fact that bacterial life is greatly influenced by the mechanics of the fluid flow both in bounded and unbounded conditions. A number of works have been conducted to understand bacterial swimming behavior near abiotic surfaces as they form biofilms near surfaces that enhances their ability to fight antibiotics ^3,24,25^. In terms of bacterial locomotion, many researchers have focused on the near wall behavior of motile bacteria ^26–29^. However, bacterial transport in flowing conditions differ from the above studies in the sense that the bacterial motion is affected by the mechanical component of fluid flow i.e., shear ^30^. In many situations, like marine and aquatic habitats, porous matrix luminal flows in human gut ^31^, bloodstream infection ^32^, etc., bacteria are driven or influenced by the external fluid transport. There are very few works concerning the adhesive behavior of bacteria near surface in shear flows. Lecuyer et al. ^33^ conducted experiments from lower to higher shear stress regime (0 – 3.5 Pa) and found that counterintuitively, in mutants with or without surface organelles, the residence time of *Pseudomonas aeruginosa (P. aeruginosa)* on glass or polydimethylsiloxane (PDMS) increased almost linearly with increase in the shear stress. Rusconi et al. ^34^ worked with motile bacteria *Bacillus subtilis* and *P. aeruginosa* at lower shear rates ranging from 0-50 s^-1^ inside microfluidic chamber and reported a cell depletion of almost 70% from low-shear regions due to getting trapped in high shear regions. Shear gradient based trapping behaviour of bacteria on non-planar surfaces was studied by Secchi et al.^35^ where they showed that bacteria prefer colonizing at the leeward side of cylindrical structure. Sherman et al.^36^ found that the microcolony of tower like structure in *Staphylococcus aureus* forms around shear stress value of ∼ 0.15 dynes/cm^2^.

Even after significant discoveries in the field of medical biology, the threat of bacteria looms over humankind. Bacterial species have been evolving rapidly that has led to the problem of antimicrobial resistance (AMR)^37–40^. Once effective medicines have started to become ineffective due to the developing resistance of bacteria against them. The acronym ‘ESKAPE’ that includes *Enterococcus faecium, Staphylococcus aureus*, *Klebsiella pneumoniae* (*KP*), *Acinetobacter baumannii*, *P. aeruginosa and Enterobacter spp* is a group of gram-positive and gram-negative nosocomial bacteria those have developed multidrug resistance (MDR) and enhanced virulence ^41^. Such evolution has resulted in declining treatment options and higher mortality rates caused by these MDR bacterial species ^42^. According to a global surveillance data ^43^, *KP* is responsible for most of the infections occurring in the hospital environments and is also the third leading bloodstream pathogen in children. It’s existence in different parts of the body like membranes surrounding brain, urinary tracts, respiratory tracts and bloodstreams, can lead to diseases like meningitis, urinary tract infection, pneumoniae, sepsis, etc.^43–46^. The presence of viable bacteria inside bloodstream is known as bacteremia^32^ that can become a life-threatening condition and result in hematogenous spreading.

*KP* is an opportunistic gram negative, rod shaped, lactose fermenting bacillus with prominent capsule. On agar media, it shows mucoid phenotype due to the presence of polysaccharide capsule attached to bacterial outer membrane^47^. Recent studies show that the prevalence of *KP* colonization ranges from 18.8% to 87.7% in Asia, and 5% to 35% in Western countries ^48^. The various virulence factors of *KP* includes capsule polysaccharide (CPS), lipopolysaccharide, siderophores and fimbriae ^49,50^. CPS is the most important virulent factor of *KP* that safeguards the bacteria from phagocytosis by polymorphonuclear granulocytes and prevents killing by bactericidal serum factors^49,51^. The O antigens in *KP* prevents it from complemented-mediated killing^49^. Siderophores can influence the host immunity by modulating cellular iron homeostasis ^52^ whereas different types of fimbriae helps in adhesion and colonization ^49^. The rise in antibiotic resistant strains are a cause of deep concern to the medical fraternity. Quinolones are the bactericidal antibiotics that are used to kill the bacteria by targeting their DNA topoisomerases^53^. *KP* is one of the most important fluroquinolones resistant pathogens ^46,54–56^ that can cause fatal diseases like bacteremia.

From the literature survey, it is evident that the effect of fluidic environment on the bacterial virulence has not been investigated. The goal of this work is to the understand how flow stresses generated purely by the flowing fluid inside micro geometries affects the bacterial population. Particularly, we try to unearth how these velocity gradients changes the morphology and virulence of the bacterial sample. Flowing sample of bacterial population is like any other metabolic activity that takes place inside a human body and such a condition can manifest in a healthy human as well. Hence, it is critical to understand the consequences of these flow stresses on the bacterial life. The experimental setup and the flow of work can be extended to complex physiological fluid like blood to study disease models like bacteremia. Here, we observe how virulence properties of bacteria changes when stressed under different conditions. To further assess the bacterial fitness, an antibiotic study with fluoroquinolone based antibiotic ciprofloxacin is conducted. This unique study investigates a largely neglected habitation of bacteria and demonstrate that *KP* shows high levels of virulence when exposed to high shear rates that are relevant to blood flows inside human body.

### Governing equations and stress on a bacterium

The rheological tests of the bacterial sample confirm the Newtonian behaviour of our working fluid (Figure S1). Figure 1 depicts the experimental setup used in the present work (details in material and methods) The choice of the *AR* (>1, here 1.6) ensures that the dominant flow gradient lies in the XY plane inside the microchannels as shown in figure 2. The 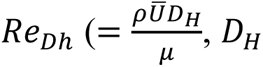 is the hydraulic diameter and *U̅* is the average velocity) values lies between 2 to 3 for all the stressing conditions. The flow condition can be modelled as a two-dimensional pressure driven Poiseuille flow between parallel plates that can be described by the continuity equation (Equation 1) and momentum equation (Equation 2) ^34,57^

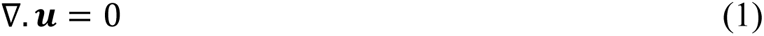

**Figure 1:**
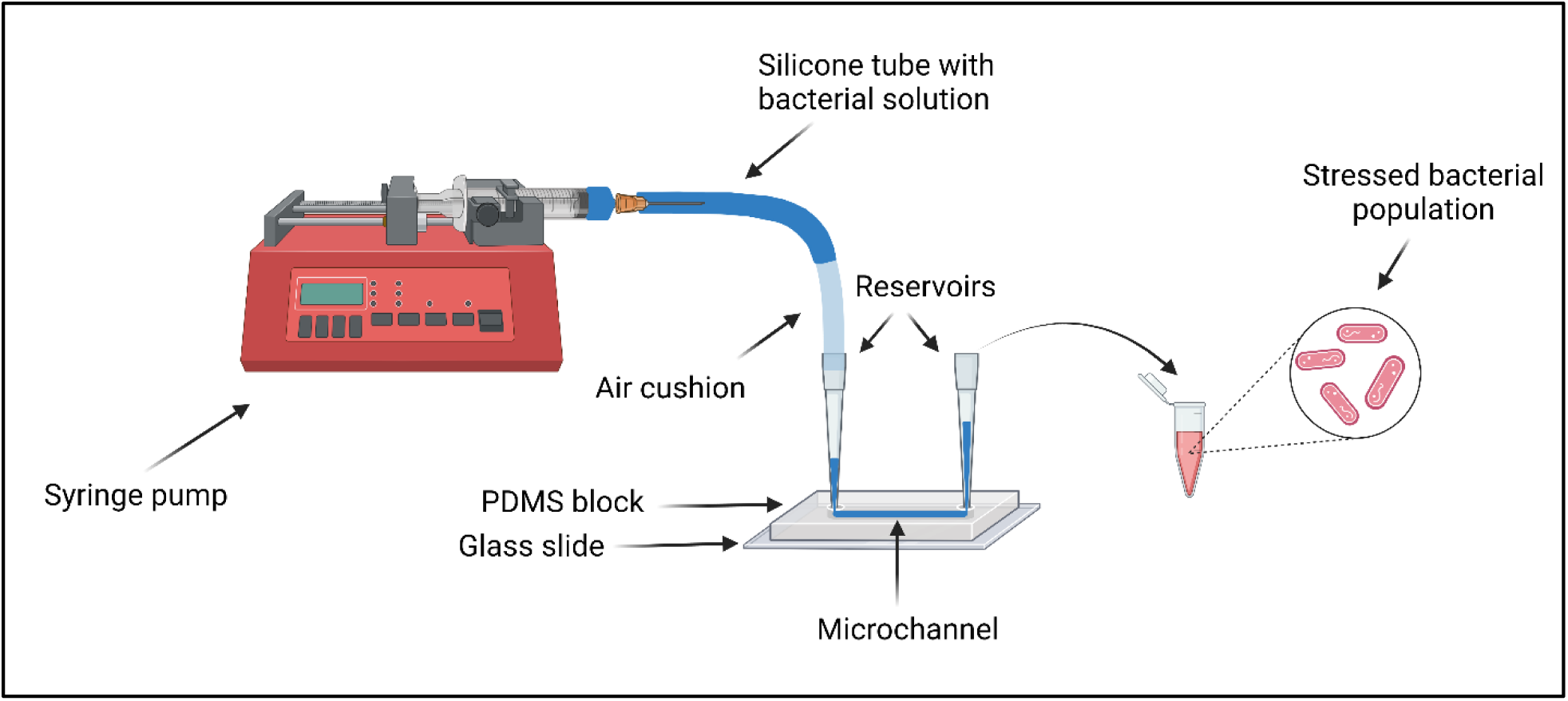
Experimental setup for stressing the bacterial sample. The dark blue colored fluid in the syringe and silicone tube represents the unstressed bacterial sample. Inside the microchannel, this fluid represents the sample being stressed and finally, in the Eppendorf tube, the red fluid represents the stressed bacterial sample. The stressed fluid is further taken to perform different assays. For more details on the experiments refer to the materials and method section.

**Figure 2:**
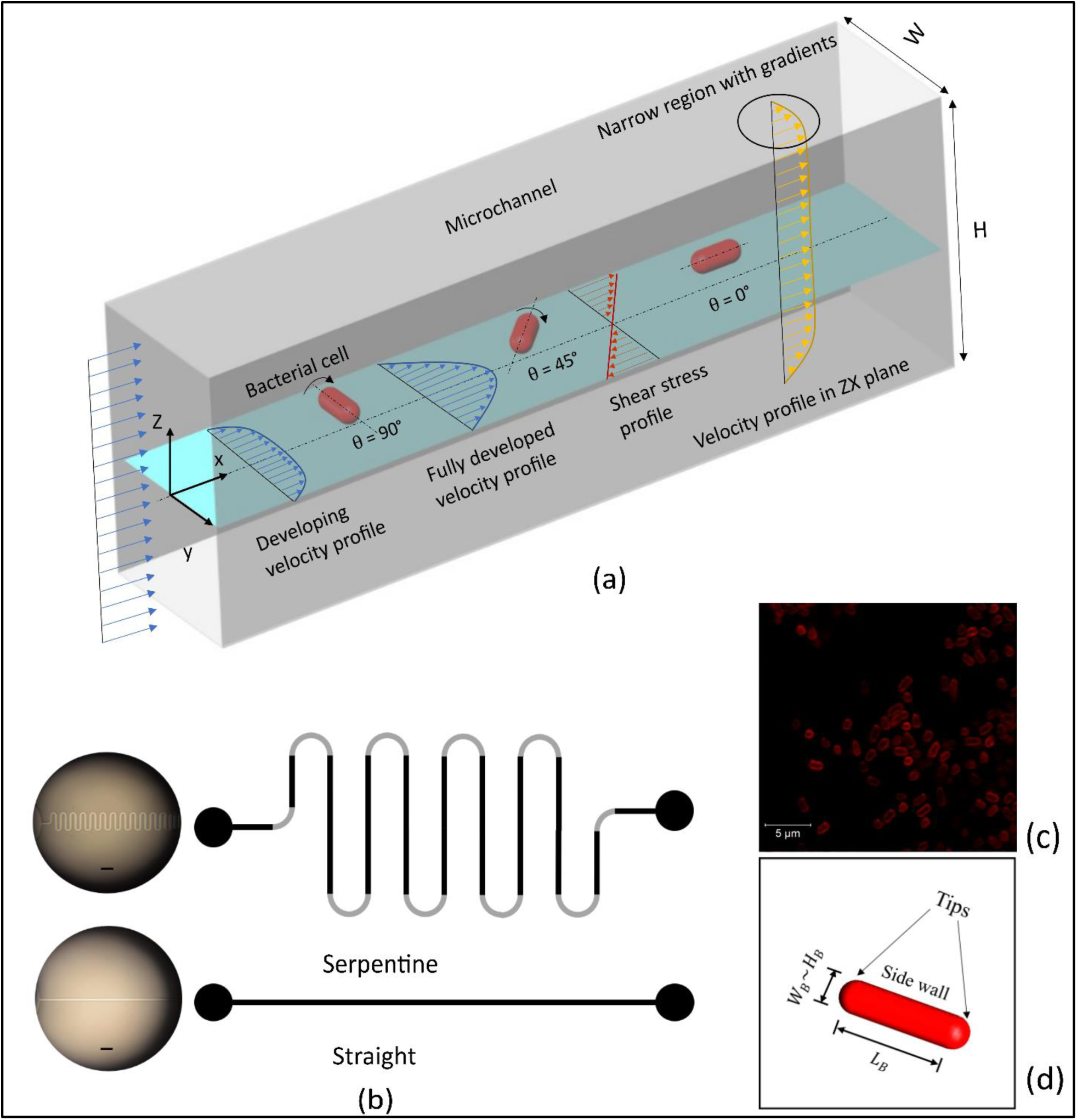
(a) The figure depicts a microchannel with *AR* > 1. Bacterial cells in XY plane at three different orientations, *θ* = 90°, 45° and 0° calculated from the centerline in an anticlockwise direction (seen perpendicular to the plane) are shown. The bacterial cells are expected to align to the flow direction (*θ* = 0°) irrespective of their initial orientation. As the fluid enters the channel, a developing velocity profile ultimately evolves into a fully developed velocity profile as shown. The velocity gradients and the shear stress profile dominate in the XY plane whereas the gradients in the ZX plane are localized only near the walls due to high *AR* of the channel (b) The two types of microchannel geometry in the present study: serpentine and straight. The images on the left are taken using bright-field microscopy. The grey area in the exaggerated schematic on the right highlights the curved part, and the black region shows the straight part which is almost 7 times more than the curved part (c) A confocal microscopic image of the bacterial sample (d) The schematic of bacteria as used in the present study for calculations.

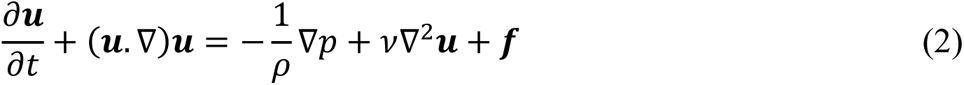

Where ***u*** is the velocity, *ρ* is the density of the fluid, *t* is the time, p is the pressure, *v* is the kinematic viscosity and ***f*** is the forcing term. The terms in bold denotes a vector quantity. The developing region or the entrance length in microchannels are generally very small ^58,59^ and depending on the present application can be safely neglected. Neglecting the forcing term (details in materials and methods) and using steady, incompressible, laminar, fully developed conditions with no slip and no penetration boundary conditions at 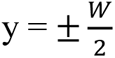, we obtain the velocity profile and shear rate equation as (equation 4 and 5 respectively):

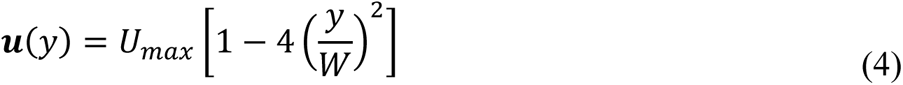

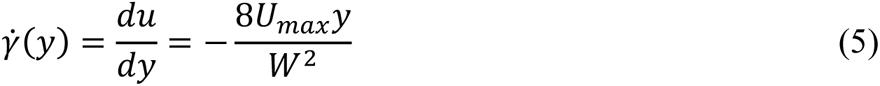

where, 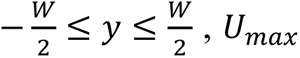 is the maximum velocity or the centreline velocity of the parabolic profile and is equal to 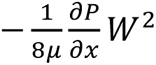. The experimental observations for velocity profiles (Figure S2) follows a parabolic profile with the maximum velocity at the centre line. The reason for existence of such a velocity profile is the no-slip boundary conditions that tends to slow down the fluid layers as we move in Z direction from wall. The shear rate varies linearly across the plane and is zero at the centre. The 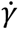 values for different stressing conditions are mentioned in Table I. The 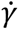 in humans can be as high as 2000 s^-1^ under normal conditions and 40000 s^-1^ under pathological conditions ^60^. The velocities inside arteries and veins can range from 0.02 cm s^-1^ - 290 cm s^-1^ ^61^. Hence, the values of 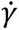 obtained in the present studies are very pertinent to flow inside blood vessels. Interestingly, the bacterial sample show high potency even under such high stress conditions making the study extremely relevant to model deadly diseases like bacteremia and other blood stream infections.

**Table I:**
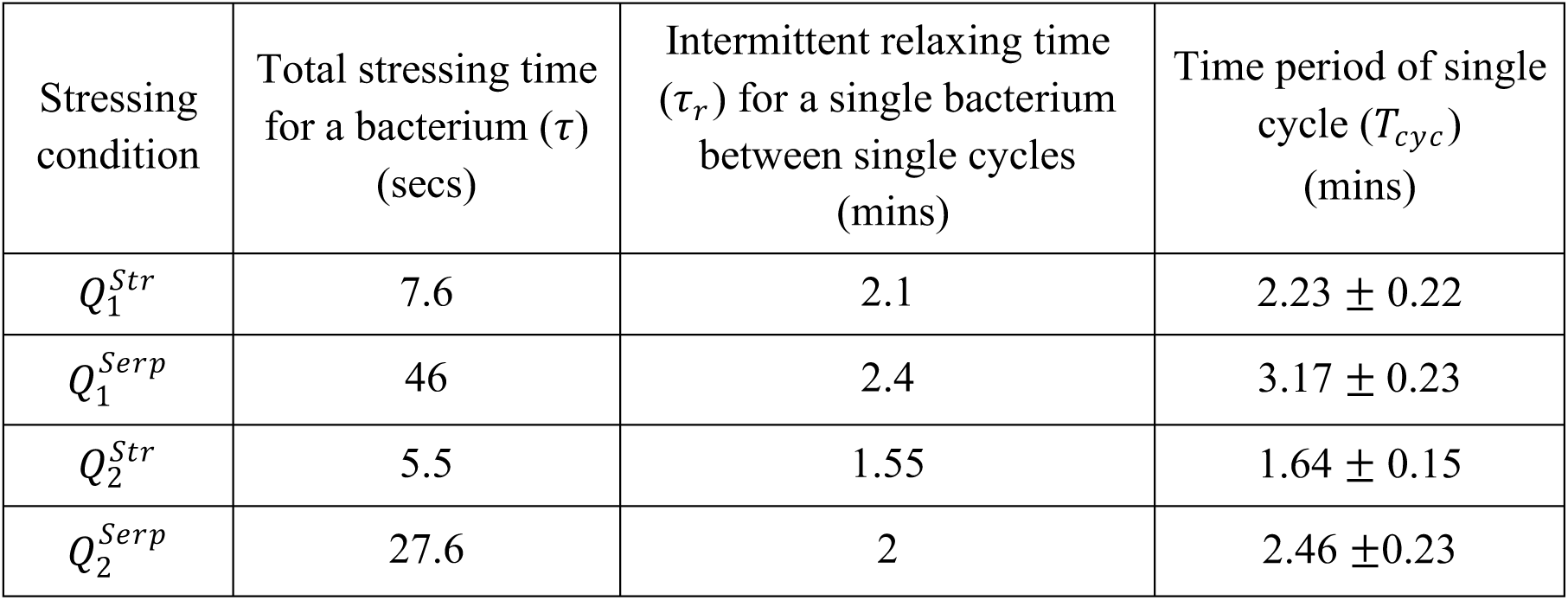
Summary of the stressing parameters used in the present study. The symbols showed for the four different stressing conditions are used throughout the paper. *Str* and *Serp* as superscript represents straight and serpentine channels respectively. The subscript *1* and *2* represents the flow rate conditions where moving from *Q*_1_ to *Q*_2_ represents increasing stress conditions. *U̅* represents the mean velocity across planes that is calculated by averaging out all the velocity values of fluid at different plane of measurements (Figure S2). 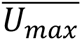 denotes the mean of maximum velocity i.e., the average of the maximum values obtained at different planes. The mean shear rate (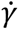) is calculated using 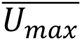 values. Higher values of 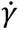 indicates higher stress values. The error represents the SD values.

*KP* is a non-motile, rod-shaped bacillus as can be seen from figure 2(c) and the authors did not find any study relating flow stresses to its virulence. It has been seen that motile bacteria are prone to shear trapping due to the local angular velocities created by the velocity gradient near the surfaces ^34,35^. However, non-motile cells simply follow the streamlines ^35^ and are distributed homogenously even in high shear regions ^7^. Majority of the previous studies considered lower shear rates and flow velocity that are at least an order smaller than what is used in the present study. The aim here is to understand the effect of such down washing of bacterial sample under constant fluid stress on the virulence of the bacterial population. It is important to note that the stresses generated due to cell-cell interaction have been completely ignored here. An important aspect of bacterial flow near surface is the formation of biofilms that helps them fight antibiotics with much more efficiency ^36,62^. On careful inspection, we found none or negligible deposition of the bacterial cells at the channel walls ruling out the possibility of forming a biofilm. This is because the flow velocity and shear rates are too high for the non-motile cells to stick to the surfaces. Furthermore, the time required for the formation of biofilm are generally higher than our experimental time (Table II).

**Table II:**
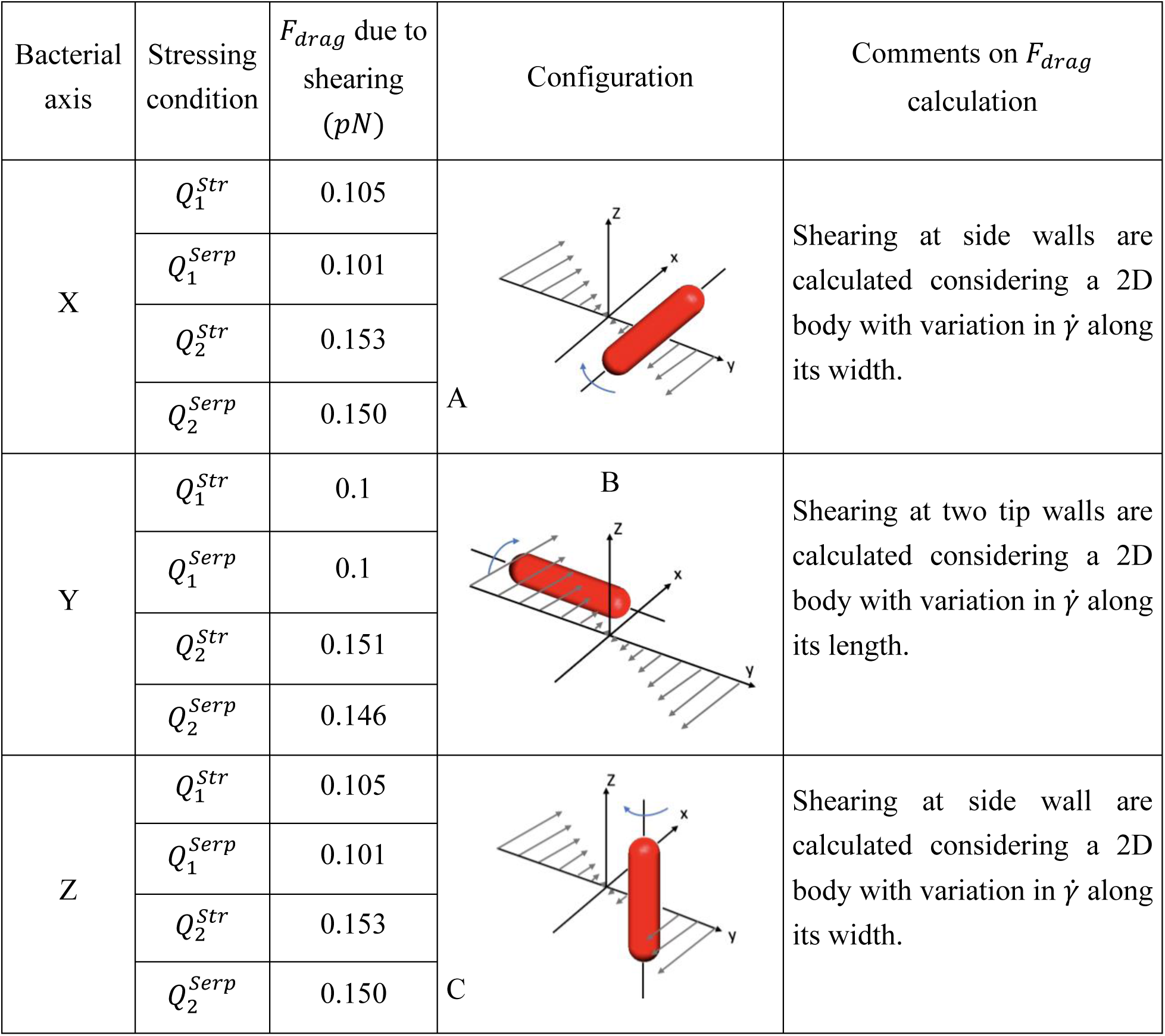
Summary of the time dependent parameters used in the present study. Total stressing time (*τ*) is calculated for one cycle as described in ‘Materials and methods’ and is multiplied by total number of cycles i.e., 20 to get the total stressing time. The intermittent relaxing time (*τ*_*r*_) is the time when a bacterial cell is expected to move out of the stressing microfluidic chamber to the reservoirs. The time-period of single cycle (*T*_*cyc*_) indicates the time taken for a single back-and-forth motion of the bacterial sample. The *T*_*cyc*_obtained large enough to ignore the oscillatory effects on the cells. The error represents the SD values.

The orientation of the bacterial cells is expected to be aligned with the flow direction. Due to experimental limitations with capturing the trajectory of a single bacterium cell at high speeds, it cannot be ascertained that if the rod shaped bacillus follows Jefferey orbit ^63^. Moreover, the bacterial population considered for the present study is high enough that hinders the process of capturing the trajectory of a single cell. The drag on each bacterium is estimated as *F*_*Drag*_ = *Aσ* = *Aμ*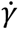 ^3^ where *A* is the exposed area of the bacteria to the flow and 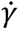 is calculated from equation 5. The general shear stress profile in the dominant plane is shown in figure 2(a). We consider that the bacterial cells are completely aligned to the flow and calculated the values of drag force for configuration A (i.e., bacterial axis along x) for different stressing condition as enumerated in Table III. However, few other plausible orientations are configurations B and C. At these configurations, the vorticity axis (if enough hydrodynamic torque is generated) will be along Y and Z direction as shown in Table III. The order of drag values obtained for both B and C configuration are same as that for A. The assumptions and methodology have been mentioned in the comments section of Table III. Moreover, due to the gradients being localised only near the wall in ZX plane, and bacteria being away from it (as considered in this discussion), the bacteria will face shear only from stress profile in XY plane that would keep on decreasing in magnitude as we go away from centreline. The normal component of drag force is expected to be order of magnitude smaller for configuration A. However, if at all the bacterial cells remain in configuration B or C, without toppling due to stress in XY planes above and below the plane shown in Table III, the normal force generated will be of similar order with slightly higher values. All the above discussion is valid only for cases when the cells do not lie exactly at the centre of the channel where the stress value becomes zero. Considering the population of cells, it can be hypothesized that the bacterial cells will be orientated at different orientations all over the channel. However, the values of drag for *Q*_1_and *Q*_2_are 0.1pN and 0.15pN respectively and the order of any kind of force on the cell remains as ∼O(0.1pN). This is a significant amount of stress from bacterial adhesion point of view as they can hold upto few to hundreds of piconewtons ^3^. The other important parameter in consideration in the present study is the value of *τ* that varies with channel geometry. Hence, the results obtained here is due to an interplay of *F*_*drag*_and *τ* values. Table II contains information about the *τ* of a single bacterium inside the microchannel.

**Table III:**
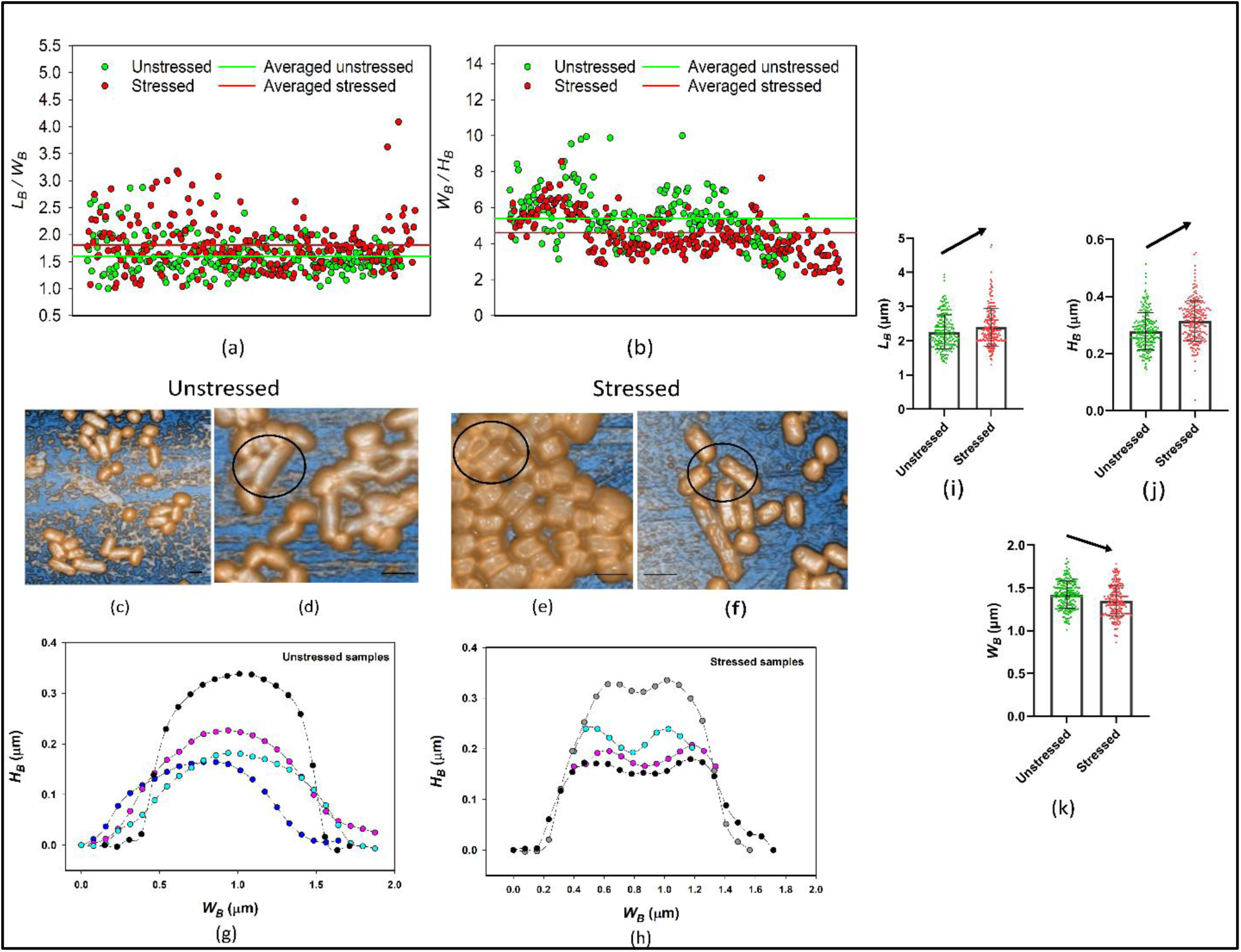
The *F*_*drag*_ at different configurations for different stressing conditions of a bacterium inside microchannel. Only three configurations are shown here as it is expected that the bacterial cells will be aligned to the flow direction. However, configurations like B and C can arise at the time of entry of a bacterium inside the microchannel which can further transit to configuration A. The probable torque direction is shown too.

### Shear stress alters the morphology of *KP*

Bacterial cells inside shear flows are subjected to continuous stretching due to the gradients in the velocity. As shown in Table III, the differential forcing at the surfaces creates a shearing action on the bacterial cell that acts like a mechanical cue for it to react. In response, we see that the morphology of the cells is significantly altered. We perform atomic force microscopy (AFM) for all the stressing conditions to understand the global trends and see how specific stress conditions modify the structural aspects of bacteria due to flow stresses. Figure 3 shows the global morphological comparison between stressed and unstressed samples. It is important to note that AFM images were taken from a dried bacterial sample. Drying itself induces stresses due to desiccation, fluid motion and starvation as observed in previous works of our group ^64,65^. Since, both stressed and unstressed sample were subjected to desiccation stresses, the effect of drying is common to both the samples and the differences in the morphology can be attributed explicitly to the flow stresses. To do so, we perform non-contact mode AFM to keep the type of stress applied to the sample unique and in line with the present aim. We see that the bacterial population responds to the flow stresses by increasing its length and height while decreasing its width. A 12.6 % increase in the *L*_*b*_⁄*W*_*b*_ ratio and a 14.2 % decrease in the *W*_*b*_⁄*H*_*b*_ ratio is observed for the stressed samples compared to unstressed samples (Figure 3(a) and (b)). These findings are unique and, to the author’s best knowledge have not been reported yet for the present stressing conditions. Furthermore, figure 3(c) and (d) shows the unstressed bacterial samples having smoother surface and edges. In the present study, majority of the unstressed sample was found to have such smooth contour whereas for stressed samples we observed dents on the top surface (Figure 3(e)) of a number of cells. The profile of these dent-like structures is plotted in figure 3(h). As can be seen in figure 3(e), these dents spanned throughout the upper surface of the bacterial body. In some cases, plasmolysis of the cells was also observed, especially for stressed samples due to hyperosmotic stress induced on the cell as can be seen from figure 3(f).

**Figure 3:**
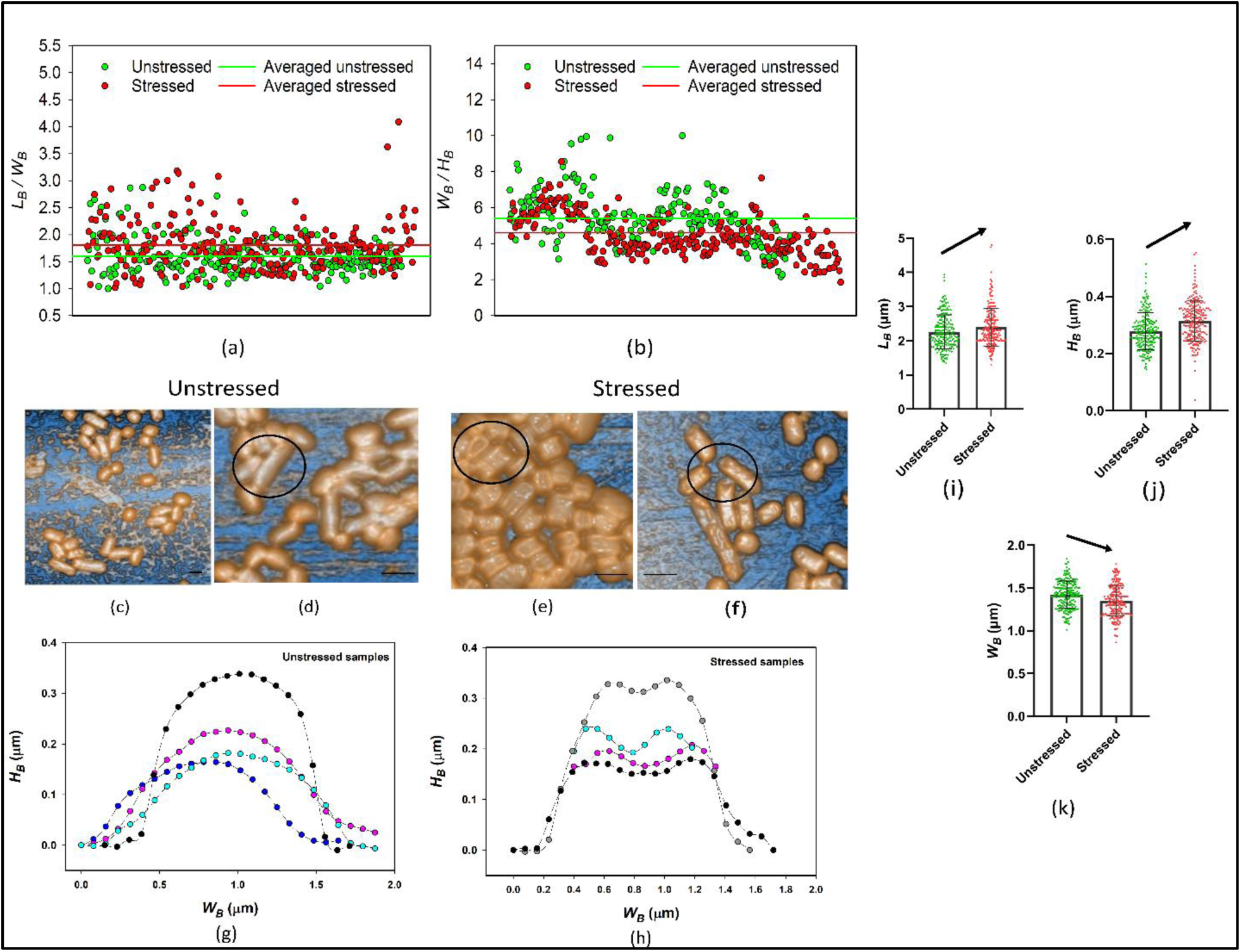
The figure depicts the global AFM observation of the bacterial cells under unstressed and stressed conditions. (a) *L*_*B*_⁄*W*_*B*_ ratio (b) *W*_*B*_⁄*H*_*B*_ ratio (c) and (d) depicts AFM images of unstressed samples. The highlighted region shows the smooth contours of unstressed samples in (d). (e) and (f) depicts AFM images for stressed samples. The highlighted region in (e) shows the dented top surface and in (f) it shows the effect of plasmolysis on the cells. (g) and (h) depicts the contour profiles of the bacterial cell for unstressed and stressed conditions respectively. (i), (j) and (k) shows the variation of *L*_*B*_, *H*_*B*_ and *W*_*B*_ for unstressed and stressed samples. The data represents the mean ± SD. All the scale bars represent 1*μm*.

Localising our analysis, figure 4 compares individual stressing conditions with corresponding unstressed situations. The maximum increase in *L*_*b*_⁄*W*_*b*_ ratio is seen in the case of 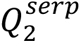 followed by 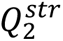 (Figure 4(d) and (b) respectively). This result depicts the importance of 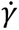 and *τ* on the bacterial morphology. The case 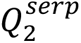 has the second largest *τ* which combined with high 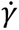 results in the maximum change whereas in cases with lower stress values i.e., cases with *Q*_1_, the changes are significantly lower. The values of 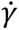 for 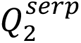 and 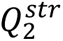 differ by a small margin (see Table I) so, the major differentiating factor between these two cases is the *τ*. However, as we hypothesise later, these bacteria can be sensitive to small amount of shear rates too and it is tough to explicitly conclude what is the exact reason behind the observed changes. This change in the dimensions of the bacterial cell is certainly a result of the mechanical strain that is sensed by the bacteria at the molecular level. There are significantly few works relating the mechanical stresses to the inner workings of the bacterial cell. Bosshart et al. ^66^ studied the effect of tensile loading on outer membrane protein A (OmpA) of *KP*. Protection from osmotic stresses by bacteria was studied by Wang et al. ^67^ where they highlighted the role of hydrated bacterial capsule in dampening the osmotic stresses. Recently, Hariharan et al. ^68^ studied the effect of impact stresses on bacterial surfaces and found significant surface damage with spikes and bumps all over the cell. The morphological changes obtained in present study are non-trivial and hence demands a deeper investigation of the bacterial strain at genetic level to understand the exact reason for such alterations.

**Figure 4:**
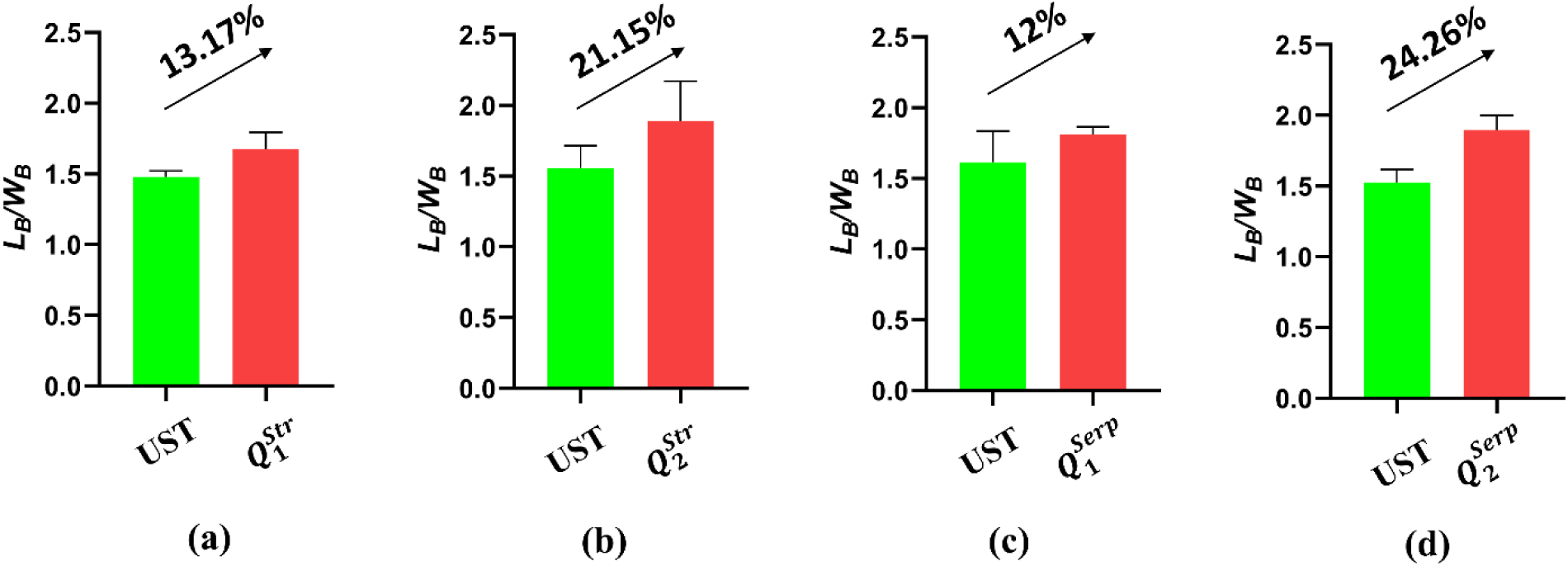
The *L*_*B*_⁄*W*_*B*_ ratio in unstressed and stressed condition obtained from AFM study. UST represents unstressed bacterial sample. The figure depicts a comparison between unstressed and stressed bacterial samples under different stressing conditions (check Table I for symbols). Here, 4 (a), (b), (c) and (d) represents each experimental set. Likewise, the test for each set was conducted more than three times and at least 200 bacterial cells were tested. The data represent the mean ± SD.

### Viability assay

Viability, unlike dead or alive, is defined as the ability of any bacteria to form progeny and is one of the most fundamental physiological state ^69^. As this area of research deals with a population of cells instead of distinctively countable number of cells, it is an extremely challenging task to assess whether the bacterial cells under stress are alive or dead. The challenge is multiplied for studies dealing with non-motile cells as motility is the best detectable sign of life. Interestingly, there can be bacteria that are alive but not culturable ^70^. However, the measure of viability is important since it provides an idea about the potential of the bacteria to undergo binary fission and form next generations of bacteria.

It has been seen that under various kinds of stress conditions like osmotic stress ^71^, thermal stress ^71^, oxidative stress ^72^, shear stress ^73^, evaporative and impact stress ^68^ results in a loss of viability of the bacterial population. We conduct different tests by keeping flowrate and geometry constant in each one of them to understand how the viability of bacterial cells changes. Figure 5(a) and (b) shows the viability plots obtained by keeping channel geometry and flow rate constant respectively. Due to experimental limitations, all the parameters could not be tested for the same sample of bacteria however, on investigation we found homogenous results for the unstressed bacterial sample confirming the quality of bacterial culture. Further, all the assays and set of experiments were carried out for at least 3 times to ensure repeatability in the results. Each set of experiment refers to an experimental condition where one of the four parameters (two channel geometry and two flow rate) is kept constant. Likewise, we perform four experiments for each of the biological tests keeping each of the parameter constant. Since, the base cases (unstressed sample) are different for each set of experiment, a cross comparison between set of experiments might lead to erroneous interpretations and hence are not discussed here.

**Figure 5:**
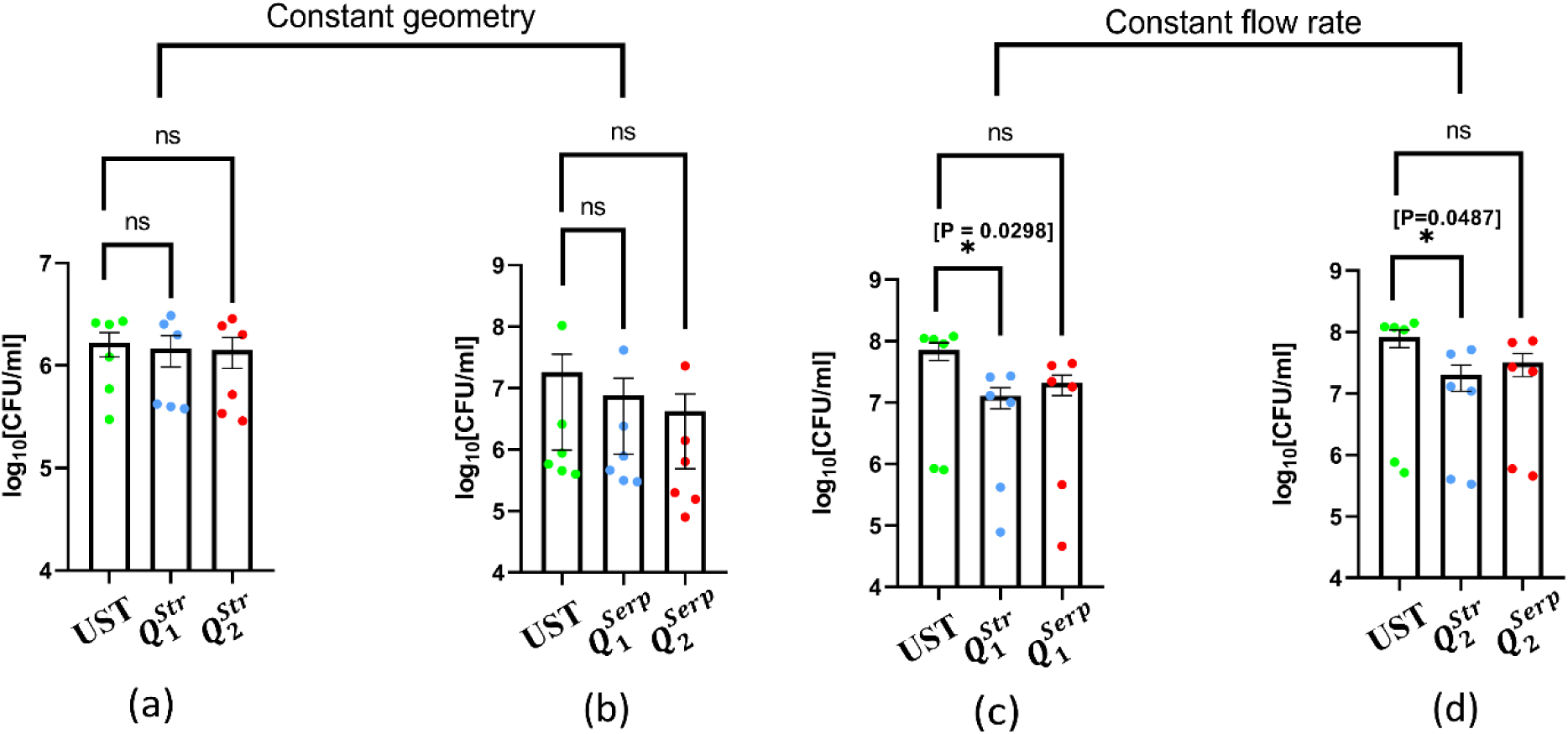
The viability of the unstressed and stressed bacterial sample are shown post 16 h of stressing. Here, UST represents the unstressed sample, and the other labels represent the different stressing condition as described in Table I. All the four figures (a), (b), (c) and (d) represent unique experimental set (*N*=5, *n* ≥3; *N* and *n* represents biological and technical replicates respectively, the average of *n* values is plotted) with different UST samples. (a) and (b) are carried out keeping the geometry constant as straight and serpentine respectively and (c) and (d) are carried out keeping the flow rate constant as *Q*_1_ and *Q*_2_respectively. The data represents the mean ± SEM. (P)* <0.05, (P)** < 0.005, (P)*** < 0.0005, (P)**** < 0.0001, ns = non-significant, Student’s t test.

A global observation from the obtained data follows that under any magnitude of flow stress, the viability of the bacterial population deteriorates. As can be seen from figure 5, this reduction is not as extreme as expected and speaks about the potency of the bacterial strain. However, the trends obtained are recurrent and is worth discussing in context of the flow stresses. As discussed above, the decrement in viability does not necessarily mean that they die during the stressing process but may also become unculturable. Hence, there is a need to develop better methodologies to estimate the amount of live and dead cells under any experimental condition. The present observation however indicates towards the inability of a certain number of bacterial cells to handle the applied stress. For both straight and serpentine channels, there is a decrease in the viability as we transit from lower to higher stress condition (Figure 5(a)). For constant flow rate experiments (Figure 5(b)) we see that for both the flow rates, viability is more in case of serpentine channels compared to straight channels although the viability decreased for both the channels when compared to the unstressed sample. As the difference between the stress values generated for straight and serpentine is not significantly different, the only other factor that may play a role here is *τ*. We hypothesize two possibilities here: ***H1*** The bacterial cells are sensitive to small changes in the stressing condition (3 – 5 %); ***H2*** Due to longer *τ* in serpentine channels (∼ 5 – 6 times compared to straight channels), the bacterial cells start adapting to the environment resulting in better handling of the applied stress and hence, exhibits slightly better viability. For ***H2***, there seems to be a possibility of an adaptive response that triggers inside the bacterial cell after being stressed for a specific time that helps them stay viable. The critical *τ* beyond which the cells start showing internal resistance and the gene expressions responsible for such a phenotypic change needs to be further studied. However, for ***H1***, a controlled parametric study needs to be carried out to understand the sensitivity of bacterial cells to the changes in shear rates.

### Intracellular proliferation of bacterial cells in RAW264.7 macrophage and phagocytosis

Virulence can be defined as ‘the relative capacity of a microorganism to cause damage in a host’ ^74^. It can also be understood as a quantitative measure of disease severity or pathogen’s fitness ^75^. In a sense, the success of pathogen depends on its ability to replicate in phagocytic macrophage cells^76^. Immune cells like neutrophils, monocytes, macrophages, etc., are present inside the tissues throughout our body in the form of white blood cells. A vital function of phagocytic cells is to perform phagocytosis that is essential for tissue homeostasis. Moreover, phagocytic cells like macrophages are called as professional phagocytes because of their ability to perform phagocytosis with high efficiency ^77^. Here, we infect the macrophages with the stressed and unstressed bacterial cells and see if they survive the killing by macrophages. We observe that stressed bacterial cells exhibits large proliferation inside RAW264.7 murine macrophages compared to unstressed cells after 16 h of infection. This behaviour of bacterial strain qualifies it to be a pathogen and holds the capacity to evade host defence mechanism. The exposure to fluidic flow stresses and how it relates to the bacterial virulence has never been reported before and forms the unique point of the present study.

Figure 6(a), (b) depicts the intracellular fold proliferation of the bacterial strain for unstressed and stressed conditions at constant geometries. The proliferation of the stressed bacterial sample is much more than that of the unstressed sample with the maximum proliferation observed in *Q*_2_ cases that have the highest 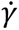. Interestingly, when we compare the effect of geometries at constant flow rates (Figure 6(c), (d)), the maximum proliferation occurs under 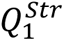 and 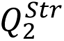 conditions compared to 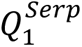 and 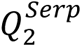 conditions respectively. The fold proliferation data can be well corroborated by the viability data where we saw the maximum depletion in the viability for 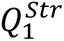 and 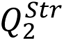 conditions. For same conditions, the intracellular proliferation observed are the highest. Using the arguments of ***H1***, this can be justified since in straight channels, the shear stress acting on the bacterial cells are slightly more, resulting in higher intracellular proliferation. This suggest that the virulence increases with increase in 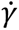 values. Whereas, from hypothesis ***H2***, we suspect that since *τ* is more in case of 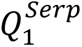 and 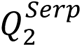, the adaptive mechanism comes into play making the bacterial cells undergo changes in their gene expressions. However, if ***H2*** were to be true, the bacterial sample should have shown more proliferation for cases with serpentine channels having larger *τ*. The results we see here are counter intuitive as the effect of *τ* seems to be less pronounced in deciding the virulence of the stressed cells as cells that are stressed for much lower time shows more proliferation. Hence, from proliferation assays, we can infer that the cases with less *τ* and more stress shows more intracellular proliferation and subsequently more virulence. This effect needs further investigation by conducting a *τ* based study to find the critical *τ* for enhancement of virulence. It is critical to note that on a global level, stressed population show much more proliferation compared to the unstressed population. This suggests that fluidic stresses turn the bacteria more virulent that can grow many folds even inside macrophage cells.

**Figure 6:**
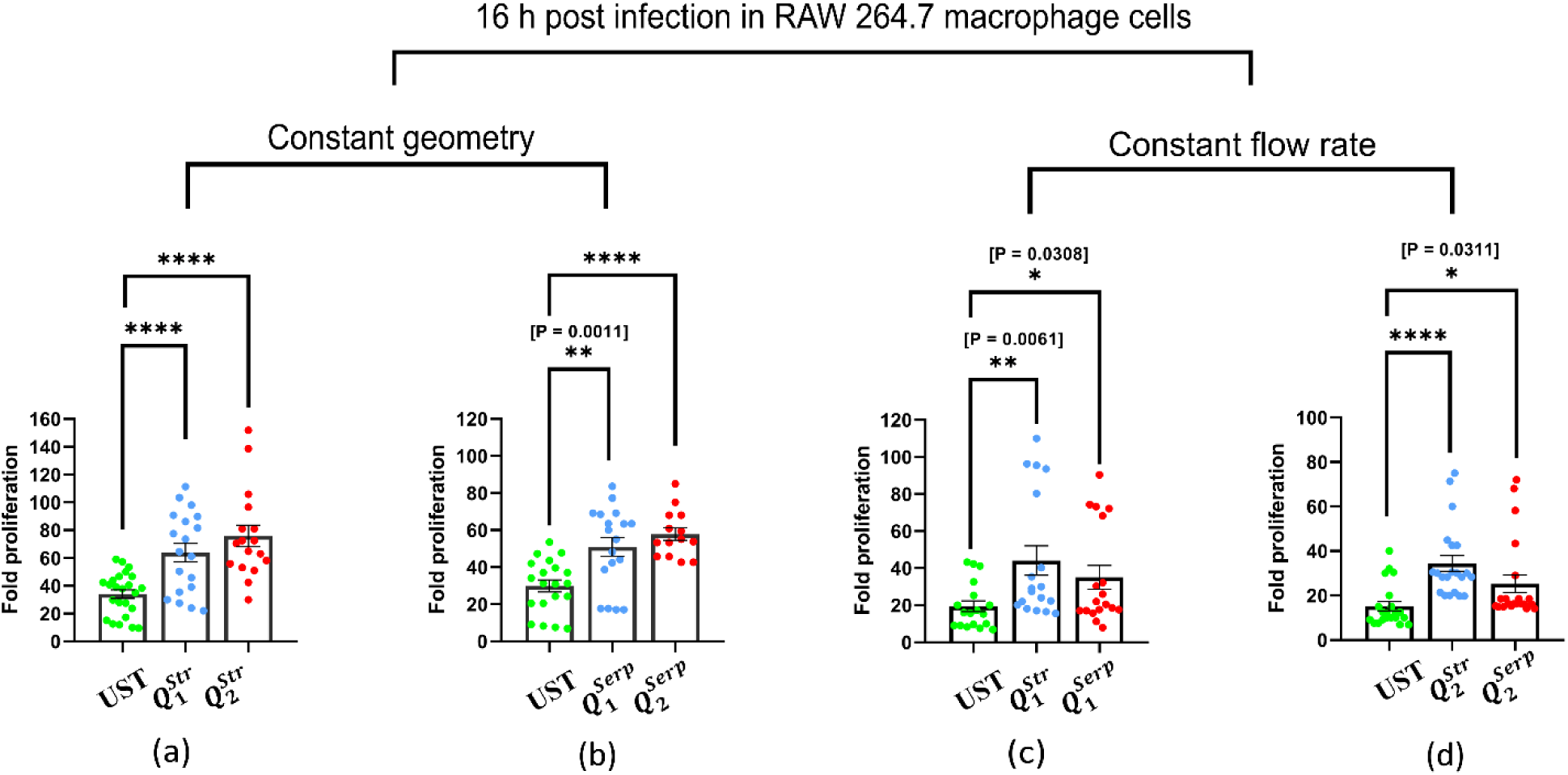
The comparison between proliferation of the unstressed and stressed bacterial sample are shown 16 h post infection in RAW 264.7 macrophage cells at various stressing conditions. Here, UST represents the unstressed sample, and the other labels represent the different stressing condition as described in Table I. All the four figures (a), (b), (c) and (d) represent unique experimental set (*N*≥3, *n* ≥3; *N*and *n* represents biological and technical replicates respectively, *n* values are plotted here) with different UST samples. (a) and (b) are carried out keeping the geometry constant as straight and serpentine respectively and (c) and (d) are carried out keeping the flow rate constant as *Q*_1_ and *Q*_2_respectively. The data represents the mean ± SEM. (P)* <0.05, (P)** < 0.005, (P)*** < 0.0005, (P)**** < 0.0001, ns = non-significant, Student’s t test.

Figure 7 shows the percent phagocytosis of the stressed and unstressed bacterial sample. Two experimental sets were carried out keeping the geometry constant and varying the flow rates. We see that the stressed bacteria are much more prone to phagocytosis by RAW 264.7 cells implying restriction in infection. The values are found to be more for maximum stress cases (*Q*_2_) suggesting the dependence of phagocytosis on the flow stresses. This difference is significant in straight channel compared to serpentine channels. Similar trends have been previously reported by our group for *Salmonella Typhimurium* subjected to mechanical stresses^68^. Ares et al. ^78^ reported the importance of cholesterol in reducing the anti-phagocytic properties of *KP* capsules. Cano et al. ^79^ suggested that *KP* manipulate phagosome maturation to survive phagocytic killing. In the context of the present study, it is important to especially assess the role CPS in protecting the bacteria from phagocytosis as highlighted by Li et al. ^49^. Different conditions can trigger different set of gene regulations or protein production resulting in such changes in the bacterial infectivity. However, it is certain that root cause of such trends obtained in figure 7 will emerge from the changes taking place at the molecular levels.

**Figure 7:**
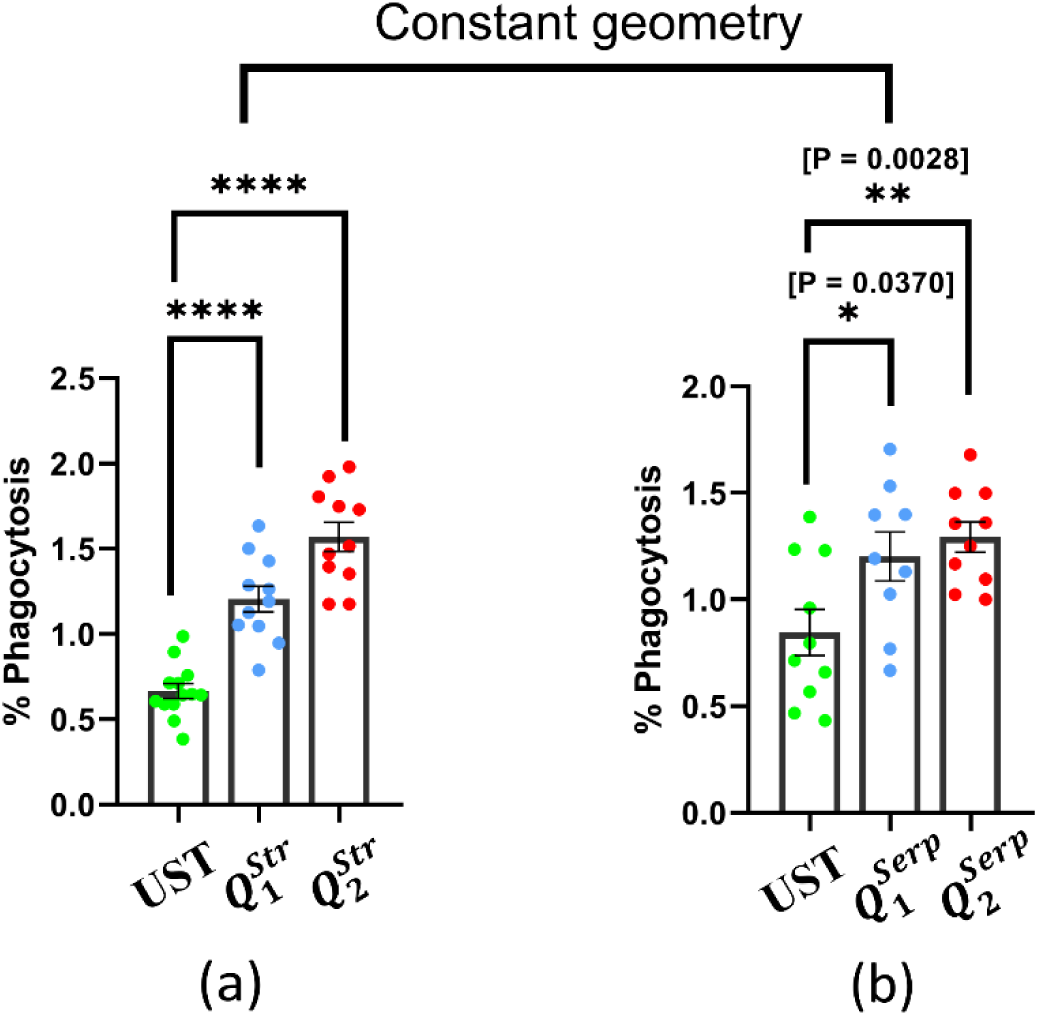
The comparison between the phagocytosis of the unstressed and stressed bacterial sample are shown by the RAW 264.7 macrophage cells at various stressing conditions. Here, UST represents the unstressed sample, and the other labels represent the different stressing condition as described in Table I. Figures (a) and (b) represent unique experimental set (*N*≥3, *n* ≥3; *N* and *n* represents biological and technical replicates respectively, *n* values are plotted here) with different UST samples. (a) and (b) are carried out keeping the geometry constant as straight and serpentine respectively. The data represents the mean ± SEM. (P)* <0.05, (P)** < 0.005, (P)*** < 0.0005, (P)**** < 0.0001, ns = non-significant, Student’s t test.

### Antibiotic survivability assays

Fluoroquinolones have been highly effective in dealing with *Enterobacteriaceae* including *KP* however, their increasing resistance against quinolones due to its widespread usage has created a problem of initial antibiotic selection against the clinical activity of *KP*. Hence, detecting and treating bacteria with adequate amount of drug has emerged as an important problem that needs to be addressed. Here, we challenge the stressed and unstressed bacterial culture at three different concentrations of ciprofloxacin (50*μ*g/ml, 100*μg*/ml and 200*μ*g/ml). The survivability is assessed by counting the CFU (colony forming unit) after plating the sample on LB agar plate (described in material and method). Figure 8 depicts the results of the antibiotic test plotted with the corresponding control (untreated sample). Compared to the control, we see a significant decrease in the viability of the bacterial sample suggesting susceptibility of the bacterial strain against ciprofloxacin. However, the count of the viable bacteria cannot be ignored for treated samples. For cases with 50*μ*g/ml and 100*μg*/ml of antibiotic in straight channels (Figure 8(a), (b), (c)), a consistent trend is observed where the minimum survivability is observed in the least stressed condition i.e., 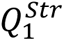. The survivability increases with increase in the stress value to *Q*_2_. This increment in the CFU suggest bacterial sample can resist antibiotic better when more stressed. At the highest challenge condition, the survivability increases for 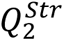. Even for serpentine channels (Figure 8(d), (e) and (f)), we observe the highest survivability in 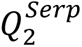 except for when the concentration of antibiotic is highest. The difference between the control and serpentine cases at 50*μ*g/ml and 100*μg*/ml is very less indicating the ability of bacteria to resist the antibiotics when stressed for longer periods. The values of CFU for 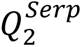 are even higher than that of unstressed samples for at 50*μ*g/ml and 100*μg*/ml (figure 8(d) and (e)). The mechanism of resistance towards antibiotic stresses and macrophage killing may be different and needs rigorous biological experimentation to be identified. Although, bacteria show susceptibility, the CFU count is high enough to highlight their AMR characteristics. The increase in CFU for more stressed bacteria indicates that under higher stress conditions, bacteria show better resistance towards antibiotics. This stressing condition however is a combination of both fluidic stresses and antibiotic stresses.

**Figure 8:**
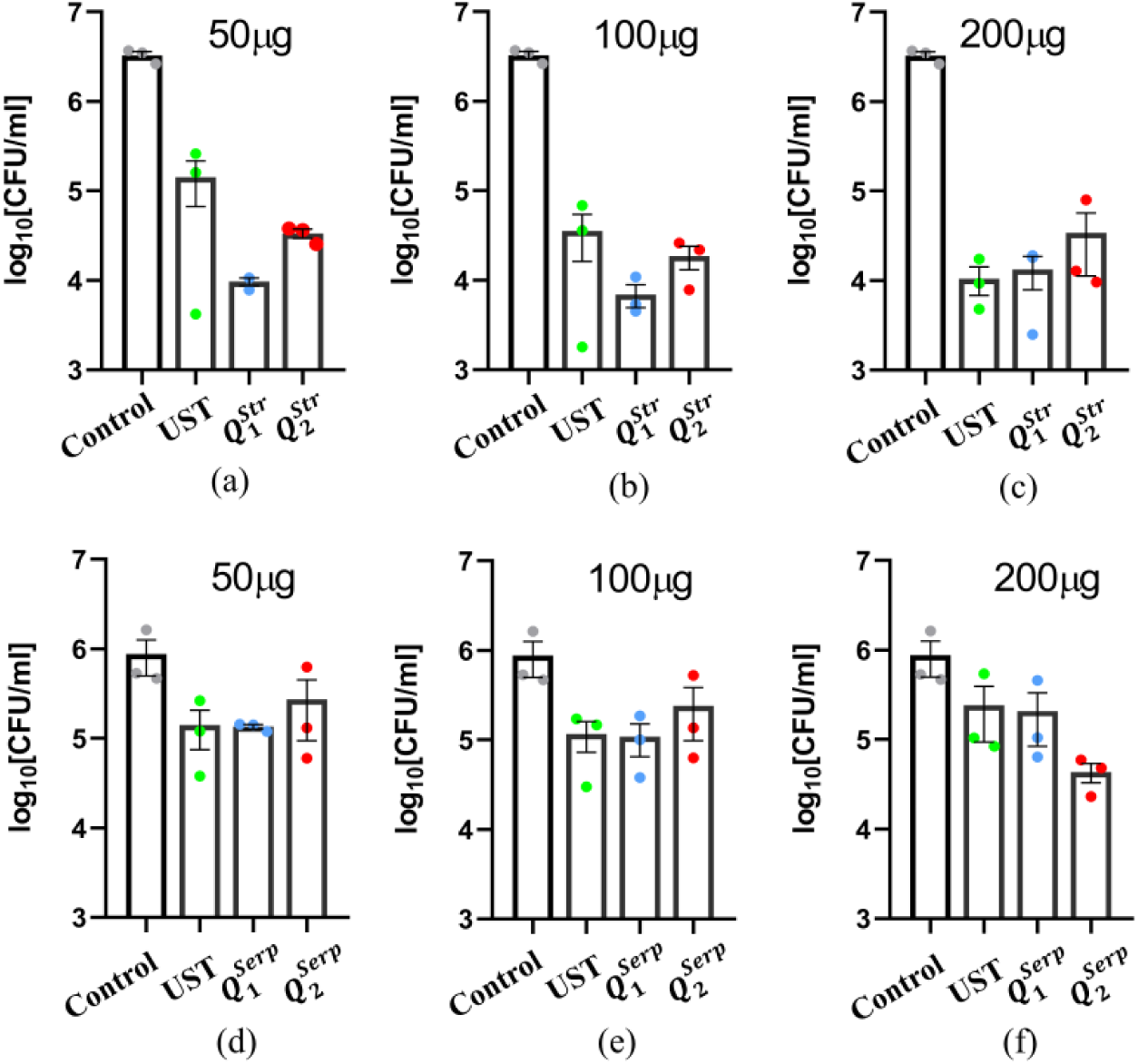
The survivability of *KP* under non-treated (control) and treated (unstressed and stressed) conditions on exposure to 50*μg*/ml, 100*μg*/ml, 200*μg*/ml of ciprofloxacin. Here, UST represents the unstressed sample, and the other labels represent the different stressing condition as described in Table I. All the six figures represent unique experimental set (*N*=3, *n* ≥3; *N*and *n* represents biological and technical replicates respectively). (a), (b) and (c) are carried out keeping the geometry as straight and (d), (e) and (f) are carried our keeping the geometry as serpentine and the flow rates are varied. The data represents the mean ± SEM.

## Discussion and conclusions

The effect of fluidic habitat on bacterial life is a largely neglected area of research. Mechanical stresses are ubiquitous to bacterial life ^3^, but very limited literature exists in this domain. To the author’s best knowledge, the relationship between fluidic stresses and bacterial virulence has never been reported before which forms the crux of the present work. This current study demonstrates how bacterial virulence changes due to fluid stresses generated by flowing fluid subjected to no-slip boundary conditions and opens new avenues to better understand the bacterial pathogenicity.

We conducted the study under four different stressing conditions by varying the flow rates and geometry of the microchannel. The choice of serpentine channels allowed us to see how *τ* affects the virulence. Under stressed conditions, the bacteria displayed significant changes in their morphology. Globally, the length and height enhance with a decrement in the width of the stressed bacteria. Locally, we see greater changes in higher stressing condition with the maximum in 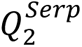. The viability of the bacterial population declined after being stressed. On comparing the effect of geometry, we find that bacterial samples show higher viability in serpentine channels compared to straight channels. We hypothesize two situations for such findings, ***H1:*** The bacterial cells are sensitive to small changes in the stressing condition (3 – 5 %); ***H2:*** Due to longer *τ* in serpentine channels (∼ 5 – 6 times compared to straight channels), the bacterial cells start adapting to the environment resulting in better handling of the applied stress and hence, exhibits better viability. The most crucial finding is the higher proliferation of the stressed bacterial cells inside RAW 264.7 macrophages compared to unstressed samples. This suggests that bacterial samples exhibit augmented virulence on being subjected to flow stresses. Furthermore, this intracellular proliferation is higher for straight channels compared to serpentine channels that corroborates with the viability findings. Hence, in general, we see that with increase in stress values bacterial samples show higher virulence. From the obtained results, we conjecture that *τ* can play a key role in triggering internal response inside bacteria and there is a possibility of existence of critical values of *τ* for different conditions. On challenging the *KP* population with ciprofloxacin at different dosages, we find that although the cells exhibit susceptibility towards the antibiotic, higher stressed cells showed much more viability compared to lesser stressed samples.

In conclusion, we infer that there exists an intricate relationship between the flow stresses and bacterial virulence that can prove to be a key to understanding bacterial pathogenicity under physiologically relevant environments. We see that stressed bacteria can demonstrate much more virulence compared to unstressed bacteria. However, the root cause of this virulence lies at the molecular level which needs to be identified by carrying out a detailed genetic analysis. The study presented here can be extended to organ-on-chip type technologies to model diseases like bacteremia and understand bacterial behaviour inside the bloodstreams.

## Materials and methods

### Microchannel fabrication

To stress the bacterial sample in a controlled way under different stress conditions, we fabricate microchannels of straight and serpentine shapes (Figure 2(b)) having 27nl and 83nl of volumetric space. The serpentine channels were made such that straight part of the channel is about 7 times more than the curved part. To ensure a dominant velocity gradient in the XY plane, we designed the channels with aspect ratio (*AR* = *H*⁄*W* = 1.6) > 1. The microfluidic devices were made using the standard soft lithography technique. Two different master moulds were prepared over a silicon chip using SU8 2050 (Kayaku advanced materials) negative photoresist. The PDMS is prepared by mixing Sylgard 184 silicone elastomer base and curing agent (Dow Corning Corporation) in 10:1 weight ratio. It is then degassed in a desiccator to remove the air bubbles and poured over the master mould before being cured for 3 hours at 60°C in a hot air oven to ensure full cross-linking of the elastomer. The inlet/outlet access holes are made using biopsy punches. Lastly, air plasma cleaner (Harrick Plasma) is used to create an irreversible bond between the PDMS chip and a glass slide that forms the base of all the channels used. The channel characterization is done using Dektak XT surface profilometer.

### Microfluidic setup for stressing the bacteria

The experimental work consists of two major part that involves stressing the bacterial sample followed by biological testing of the stressed and unstressed samples. For generating a pressure driven flow, a syringe pump (New Era pump systems) is used in pumping and withdrawal modes. As shown in figure 1, the syringe (1 ml) on the syringe pump is connected to the microchannels through a silicone tube (2mm ID) and microtip (Tarsons). Around 80% of the silicon tube is filled with the bacterial sample and the rest is left sample free to provide an air cushion to prevent the mixing of the stressed sample with the unstressed bacterial sample inside the silicone tube. A small sample of bacterial population is then sucked in through the microtip which is tightly fitted to the microchannel inlet and outlet holes making a continuous fluidic system as shown (Figure 1). The microtips at both the ends acts as reservoirs for the sample and is at the atmospheric pressure at the outlet port. The syringe pump is made to work repeatedly at pumping and withdrawal mode making the sample pass through the microfluidic channel 40 times i.e., 20 cycles (∼ 200mm). The time-period (*T*_*cyc*_) of the back-and-forth motion of the bacterial sample is very large or the frequency is very small (check Table II) and hence, the oscillatory forcing for pressure term in the NS equation is not considered. We employ two flow rates in two different geometries that results in four stressing conditions as enumerated in Table I. Once the stressing is done, the bacterial sample is collected in an Eppendorf tube and are subjected to biological testing within 30 - 45 minutes. The motivation behind choosing a serpentine geometry with much longer straight portion compared to the curved portion is to increase the stressing time (*τ*) on the bacterial sample as can be seen from the values obtained (Table II). The velocities obtained inside the serpentine geometry is slightly lesser than what is obtained in straight channels and hence less shear stress is generated inside serpentine channels at a given flow rate. It is important to note that this difference in the shear values generated for straight and serpentine channels vary only by about 3 – 5% However, overall experimental time increases significantly (42.3% for *Q*_1_ and 50% for *Q*_2_) for serpentine geometry due to the increase in the surface area of the channel. Furthermore, on changing the flowrate from *Q*_1_ to *Q*_2_ the shear rate value increases by 52% and 55 % for straight and serpentine channels respectively. The bacterial particle *Re* (∼10^−3^; = *Re* ∗ (*a*_*p*_⁄*D*_*h*_)^2^ ^16^ where *a_p_* is the characteristic length of a *KP* bacterium cell) and the Stokes number (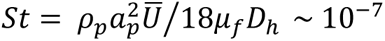 << 1 where subscript *p* and *f* represents particle and fluid respectively, *U̅* is the average velocity (Table I)) being very small, the bacterial cells are expected to follow the flow faithfully without exhibiting any inertial effects. The Peclet number (*Pe* = *U̅L*⁄*D*_*B*_, *L* is the channel length and *D*_*B*_ diffusivity of *KP* ^80^) value remains ≫1 indicating a purely advection driven flow. Hence, *τ* is calculated as the actual distance covered by a single bacterium divided by the average velocity of the fluid (Table I and II). The *τ* values are calculated with the assumption that there is no mixing in the bacteria fluid in the reservoir. After every half cycle, the fluid reaches the other end of the channel, and the cycle is then reversed. During this period, it is expected that there are no or negligible fluid stresses on the bacterial cells as they are stored in the reservoirs having diameter in millimetre in stagnant condition.

### Experimental conditions

All the experiments were performed at room temperature (25-27°*C*) and a relative humidity of 45-50 %. The stressing of the bacterial sample was carried out during their stationary phase (Figure S3) when they are not expected to reproduce. Each microchannels were used for a maximum of three times before being replaced by a new one. Before each experiment, the repeated channels were properly cleaned, heated, and inspected to make sure no bacterial colonies remained from the previous experiments. No other consumables were reused for the experiments.

### Particle tracking and imaging inside microchannel

The velocity profiles inside the microchannels are obtained by tracking neutrally buoyant red fluorescent particles (Fluro-Max, size: 860 ± 5 nm, procured from Sigma-Aldrich) added to the base fluid in an appropriate amount. A high intensity laser source (Cobalt 04 series, 532nm, 300mWatt) was utilized to illuminate the fluorescent particles through the use of dichroic filters as shown in Figure S4 that was made to fall on a di-choric mirror. The emitted fluorescent signatures were captured using a high-speed camera (FASTCAM Mini UX 100) at 10000-12000 Hz. The recorded images were then processed in ImageJ software where multiple particles were tracked at different locations inside the channel. The experiments were repeated at least 3 times to ensure accuracy and repeatability of the flow pattern inside. The velocity profiles obtained (Figure S2) is a result of the average between both the pumping and withdrawal modes as no significant difference was observed in working under these two modes.

### Atomic force microscopy of bacterial sample

The stressed and unstressed samples are made to dry over a plain glass slide under normal atmospheric conditions (Temperature: 25-27°C and relative humidity: 45-50 %.). The dried precipitate patterns are viewed under a bio-Atomic Force Microscope (Park System, South Korea) that is integrated with an optical microscope with an X-Y flat scanner to observe the physiology of bacteria. We use this scanner in non-contact mode (not to further stress the bacteria) only to acquire microscopic images. The XEI, XEP software integrated with the instrument is used to analyse the images and to estimate various mechanical parameters.

### Preparation of bacterial suspension

*Klebsiella pneumoniae* (*KP*) was cultured in Luria Bertani (LB) broth under shaking conditions at 37°C. All experiments employ overnight cultures inoculated from a single colony from the freshly streaked plate. 1 ml of the overnight cultures was centrifuged at 6,000 RPM to pellet the bacterial cells and then washed with autoclaved MilliQ water. The resultant pellet was then resuspended in MilliQ water and adjusted to the bacterial concentration of 10^9^ CFU/mL.

### Viability and infection assay

To test the viability of the bacteria under stressed conditions, the suspension was serially diluted before being plated onto an LB agar plate. To calculate the viability of stressed bacteria, CFU was divided by the dilution factor to express the result in CFU/ml. As previously mentioned, murine macrophage RAW264.7 cells were used to test the infectivity of the bacteria. RAW264.7 cells were seeded on 24 well plates and infections were given with stressed or unstressed bacteria. In addition, 24 well plates are centrifuged at 500–700 RPM to enhance bacterial adherence to host cells and incubated for 25 min at 37°C and 5% CO2. The bacteria-containing media was discarded and washed three times with 1X phosphate buffer saline (PBS). The cells are then treated with Gentamicin dissolved in DMEM for one hour at a dosage of 100μg/ml to get rid of extracellular bacteria. The cells are kept in DMEM with 25μg/ml gentamicin throughout the experiment. At 2 hours and 16 hours after infection, infected cells are lysed using 0.1% Triton-X 100, and the appropriate dilutions are spread on LB agar plates. Fold proliferation was calculated as CFU at 16 hours divided by the CFU at 2 hours. Percentage phagocytosis was enumerated by dividing CFU at 2h by CFU at Pre-Inoculum and multiplying by 100.

### Survival assay

*Klebsiella pneumoniae* was inoculated in LB broth and allowed to grow overnight at 37°C. 500µl overnight culture was pelleted down at 6000rpm for 5 mins and washed with autoclaved MilliQ water. Then pellet was resuspended in sterile MilliQ water and adjusted the bacterial number to 10^9^ CFU/ml. Stressed and unstressed bacteria were challenged with Ciprofloxacin (50μg/mL, 100μg/mL, and 200μg/mL) prepared by microbroth dilution method using MH broth and incubated for 2 hours. The CFU of the stressed and unstressed culture was enumerated after plating onto LB agar plate.

### Statistics and reproducibility

The particle tracking experiments inside the microchannel for finding the velocity profiles and shear rate values has been repeated at least three times with 10 particles tracked at the same *Z*^∗^value. The rheometer experiment for evaluating the viscosity of the bacterial sample is conducted three times. Every biological experiment has been repeated multiple times to maintain accuracy as has been indicated in the figure captions (n = technical replicates and N = biological replicates). Wherever possible and as mentioned in the respective figure captions, the unpaired students t-test is performed using GraphPad Prism 8.4.3 software. The mean with standard error mean (SEM) and standard deviation (SD) was also determined with the help of GraphPad Prism 8.4.3 software.

## Supplementary data

Supplementary sheet is attached containing Figures S1, S2, S3 and S4.

## Supporting information

Supplementary sheet

## Acknowledgment

We thank the electron microscopy facility of IISc, AFMM facility of IISc, Atomic force microscopy facility of BSSE, IISc and confocal microscopy facility and real-time facility of Dept. of Microbiology and Cell Biology, IISc.

## Author contributions

Conceptualization: S.B. and D.C. Methodology: S.B., D.C., S.J. and A.S. Investigation: S.J., A.S., N.T., A.N., R.C. Visualization: S.J., N.T. Funding acquisition: D.C. and S.B. Project administration: D.C. and S.B. Supervision: D.C. and S.B. Writing— original draft: S.J. Writing: editing and revision: S.B., D.C., A.S., N.T.

