## Supplementary sheet for "*Klebsiella* Pneumoniae turns more virulent under flow stresses in capillary like microchannels"

### SUPPLEMENTARY DATA

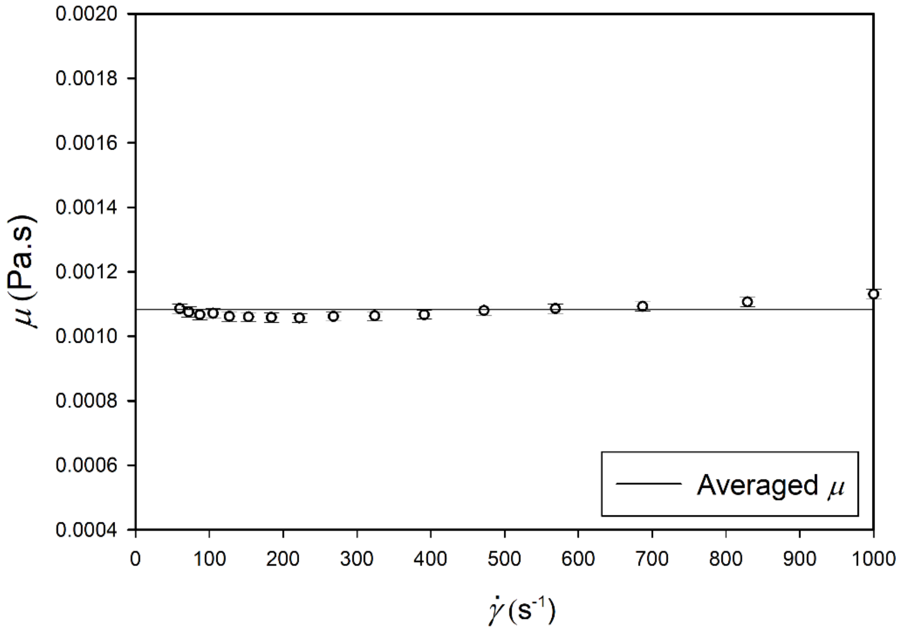


Figure S1: $Viscosity$ ($\mu$) vs shear rate ($\dot{\gamma}$) curve for bacterial solution shows a constant viscosity value with increase in the shear rate. The graph plotted is a result of three experimental runs and the error bars depicts the SD values.

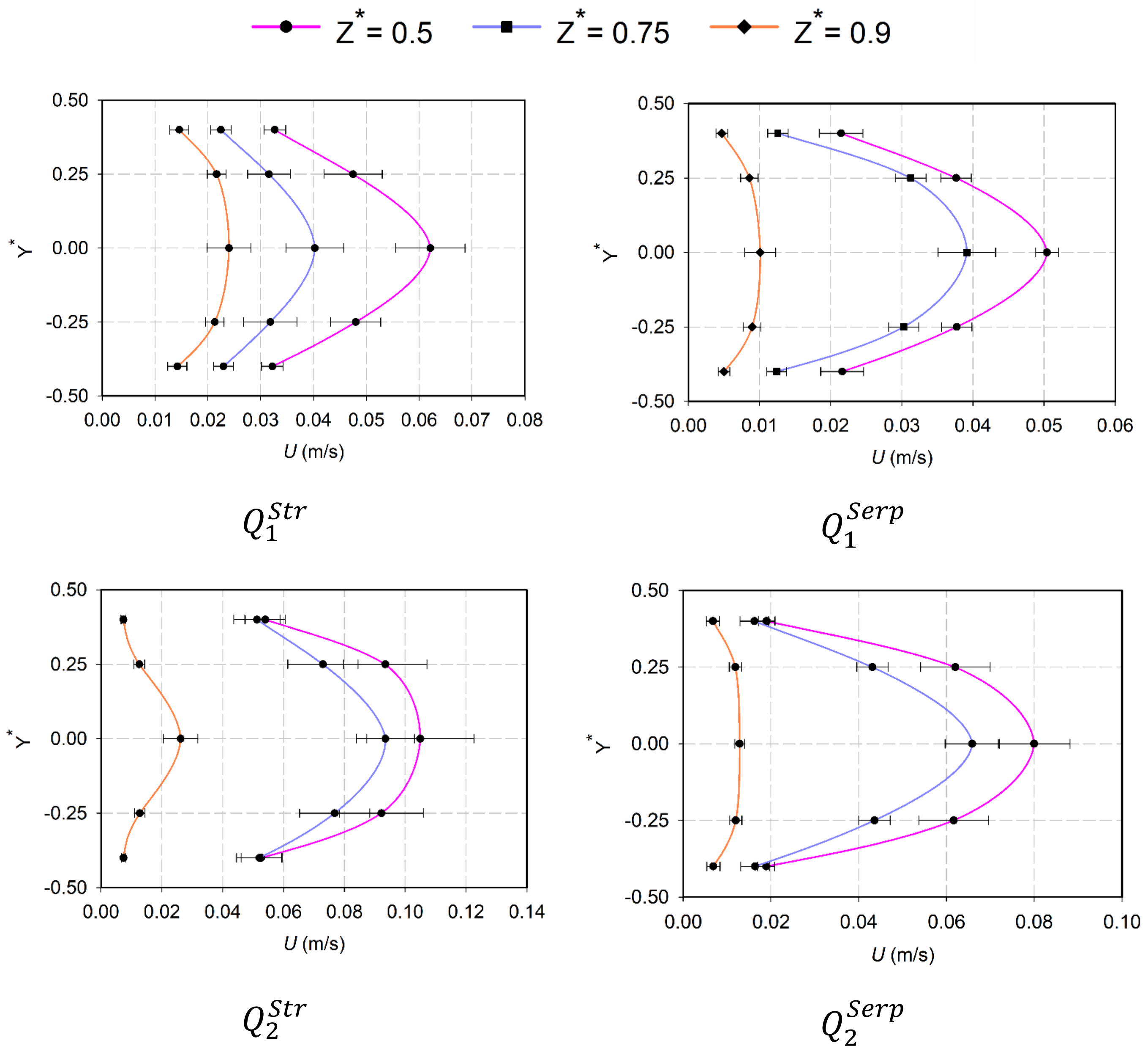


Figure S2: The velocity profiles obtained experimentally at different planes are shown for different stressing conditions. Here, $Y^* = Y/W$ represents various points in the XY plane where the measurement has been made and $Z^* = Z/H$ represents the XY planes in the Z direction. The error bar denotes the SD.

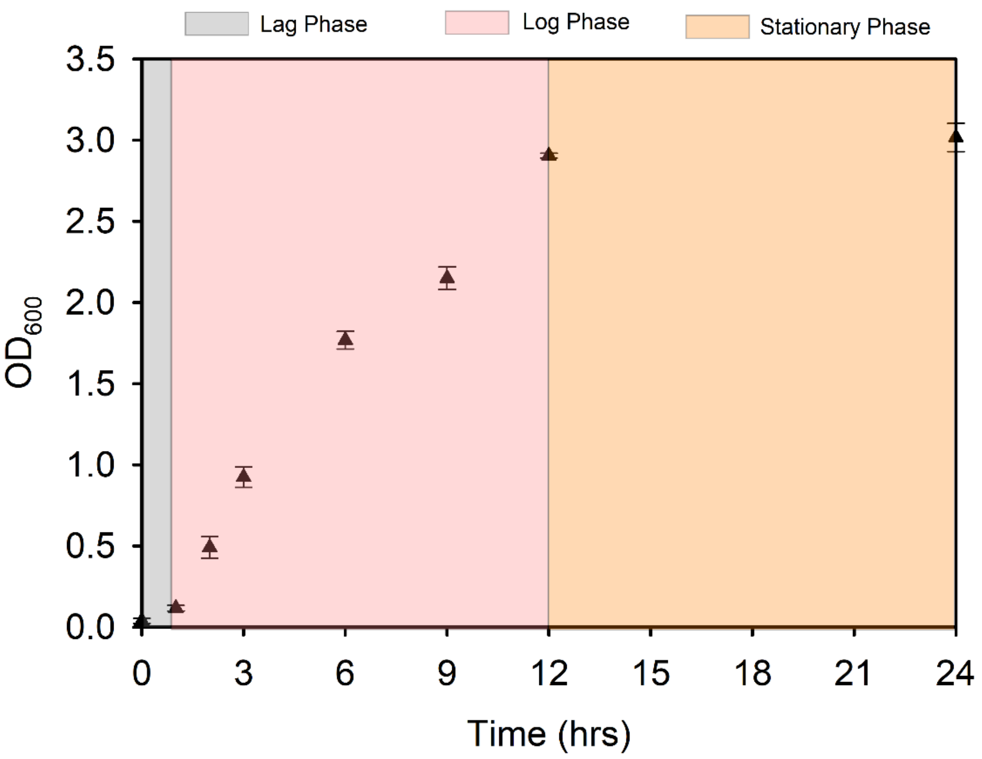


Figure S3: Optical density (OD at 600 nm) vs time (hrs) for the bacterial sample used in the present study. The three phases of bacterial growth are shown for 24 hrs. All the experiments conducted during this work were carried out in the stationary phase of the bacterial sample. The error bars depict the SD.

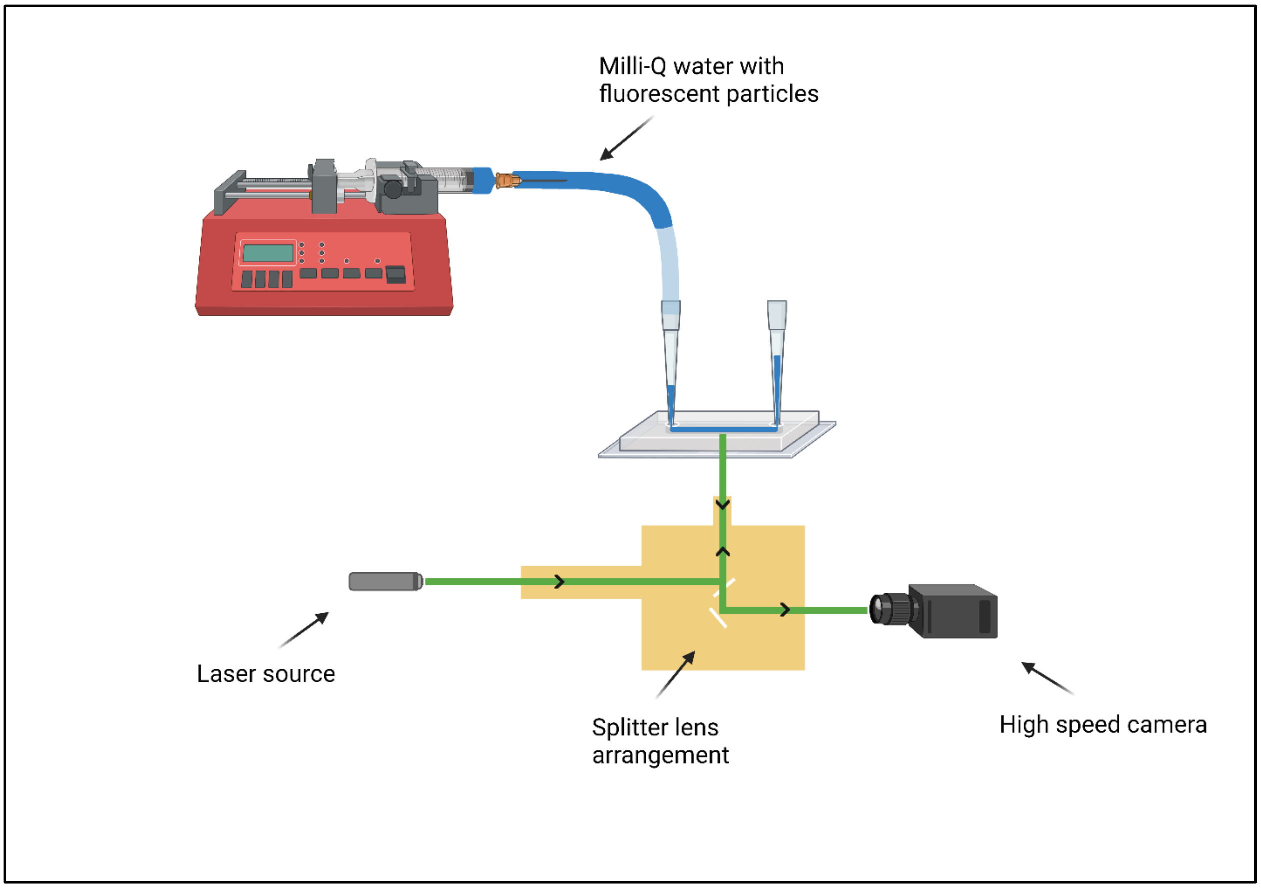


Figure S4: The experimental setup for measuring the velocity profiles inside the microchannel. A splitter lens arrangement is employed to selectively return the fluorescent signals from the fluorescent particles on getting excited by laser light (532nm). The images are captured using high speed camera at 10000 to 12000 Hz. The plots obtained are shown in figure S2.
